# Wnt signaling and contact-mediated repulsion shape sensory dendritic fields

**DOI:** 10.1101/2023.09.14.557812

**Authors:** Christopher P. Tzeng, Kang Shen

**Affiliations:** Howard Hughes Medical Institute, Department of Biology, Stanford University, Stanford, CA 94305

**Keywords:** Wnt, Frizzled, sensory neuron, field size, retraction, dendritic tiling

## Abstract

The complete and non-redundant coverage of sensory tissues by neighboring neurons enables effective detection of stimuli in the environment. How the neurites of adjacent neurons establish their boundaries to achieve this completeness in coverage remains incompletely understood. Here, we use distinct fluorescent reporters to study two neighboring sensory neurons with complex dendritic arbors, FLP and PVD, in *C. elegans*. We quantify the sizes of their dendritic fields, and identify CWN-2/Wnt and LIN-17/Frizzled as a ligand and receptor that regulate the relative dendritic field sizes of these two neurons. Loss of either *cwn-2* or *lin-17* results in complementary changes in the size of the dendritic fields of both neurons; the FLP arbor expands, while that of PVD shrinks. Using an endogenous knock-in mNeonGreen-CWN-2/Wnt, we find that CWN-2/Wnt is localized along the path of growing FLP dendrites. Dynamic imaging shows a significant braking of FLP dendrite growth upon CWN-2/Wnt contact. We find that LIN-17/Frizzled functions cell-autonomously in FLP to limit dendritic field size and propose that PVD fills the space left by FLP through contact-induced retraction. Our results reveal that interactions of dendrites with adjacent dendrites and with environmental cues both shape the boundaries of neighboring dendritic fields.

**Highlights:**

▫ Secreted Wnt CWN-2 and cell-autonomous activity of neuronal LIN-17/Frizzled receptors restrict FLP dendritic field sizes
▫ Endogenously tagged CWN-2/Wnt is punctate and visible in the same plane of growing FLP dendrites
▫ Growth of developing FLP dendrites is inhibited upon contact with extracellular CWN-2/Wnt and with PVD dendrites

## Introduction

For efficient sensory perception, neurons must be positioned appropriately to detect stimuli from the environment. For functionally similar neurons within a given subtype, their axons and dendrites completely innervate a given area of sensory tissue so that gaps in coverage are minimal, and the neurites of neighboring neurons segregate to avoid overlap and redundancy. These two criteria in like-type sensory neuron patterning have been demonstrated throughout the animal kingdom. For instance, studies in fly da sensory neurons (Emoto et al., 2004; Grueber et al., 2002, 2003; Sugimura et al., 2003), vertebrate retinal ganglion cells (Perry and Linden, 1982; Wässle et al., 1981), zebrafish trigeminal ganglia (Sagasti et al., 2005), and leech nociceptive neurons (Blackshaw et al., 1982) have established that growing neurites from neighboring neurons repel from one another following contact. Such contact-dependent repulsion is crucial for the proper placement of adjacent axons and dendrites.

Mechanosensory neurons that underlie our sense of touch also tile the skin and body wall of many organisms (Albeg et al., 2011; Blackshaw et al., 1982; Grueber and Sagasti, 2010; Grueber et al., 2003; Kuehn et al., 2019; Sagasti et al., 2005). In the nematode *Caenorhabditis elegans (C. elegans)*, harsh touch, sound, temperature, as well as other nociceptive and proprioceptive stimuli are in part detected by two sensory neurons, PVD and FLP (Chatzigeorgiou et al., 2010; Goodman and Sengupta, 2019; Iliff et al., 2021; Tao et al., 2019), with elaborate dendritic arbors whose higher order dendrites innervate the space between the muscle and hypodermis. Their collective dendrites orthogonally branch to form repetitive menorah-like structures that span the entire body of the worm with minimal gaps or points of crossing. Previous reports have documented such dendritic tiling between PVD and FLP neurons (Albeg et al., 2011; Yip and Heiman, 2016), but the mechanisms that set the boundaries of their receptive fields have remained unsettled.

Here, we find that both cell-cell contact-dependent and -independent mechanisms result in the normal dendritic field sizes for FLP and PVD neurons. We leveraged recently published single-cell RNA sequencing (Taylor et al., 2021) to discover a novel gene, *Y48G10A.6*, whose expression is specific to only PVD and FLP neurons, and use its promoter in an intersectional strategy to robustly label FLP and PVD with distinct fluorescent reporters. Dynamic imaging of this dual-color strain revealed clear points of contact and retraction between neighboring FLP and PVD dendrites. We additionally find that the secreted Wnt glycoprotein *cwn-2*, homologous to vertebrate *Wnt5*, and its Frizzled receptor *lin-17* restrict the dendritic field size of FLP. Loss of function mutants of either gene result in an expansion of FLP dendritic fields and a complementary reduction in PVD dendritic field size. An endogenous fluorescent tag of *cwn-2/Wnt* showed that forward growth of FLP dendrites is impeded upon contact of the dendrite with a *CWN-2/Wnt* labeled punctum. We find a cell-specific role for the LIN-17/Frizzled receptor in FLP, but not PVD, to restrict dendritic field size. These data demonstrate that contact-induced repulsion of neighboring neurites, as well as secreted extracellular cues like CWN-2/Wnt, both coordinately function to determine the field sizes of adjacent, within-class sensory neurons.

## Results

### Visualization of FLP and PVD morphology

Because neuronal tiling has been documented between neurons of the same functional type, the unambiguous visualization of dendrites or axons from adjacent morphologically complex neurons has been challenging. Studies (Albeg et al., 2011; Iliff et al., 2021; Yip and Heiman, 2016) have documented the tiling of FLP and PVD neuron dendrites in *C. elegans* using transgenic reporters where FLP, PVD, and other touch-sensitive neurons were all fluorescently labeled. However, the labeling of additional neurons in this strategy can obscure the complex morphology of both neurons. Cell-specific promoters that differentially label FLP versus PVD neurons are needed to study tiling and dendritic fields in this system.

To generate these tools, we leveraged CeNGEN, a single-cell RNA sequencing database (Taylor et al., 2021), to curate a list of 25 putative promoters with specific expression in either FLP or PVD (**Supplementary Table 1**). We cloned 25 promoter-fusion constructs to drive RFP or GFP expression and created transgenic arrays of each construct through microinjection. Of the resulting strains generated, 5 exhibited fluorescence in PVD and/or FLP (p*F15G9.1::mScarlet*, p*shw-3::GFPnovo2*, p*T03F6.4::GFPnovo2*, p*valv-1::mScarlet*, and p*Y48G10A.6::mScarlet*). Four of these five strains exhibited bright GFP or RFP fluorescence in other cells, and none exhibited cell-specific expression in only PVD or FLP.

However, one of these promoter fusions (p*Y48G10A.6::mScarlet*) exhibited RFP expression specifically in both PVD and FLP neurons, highlighting their body-long dendritic fields (**Figure 1A**). As the *ser-2* promoter is expressed in PVD but not in FLP (Tsalik et al., 2003), we designed an intersectional ZIF-1 based strategy (Armenti et al., 2014) to degrade *pY48G10A.6*-driven RFP and express GFP in PVD neurons (**Figure 1B**). This strategy achieved cell-specific expression of RFP in FLP and GFP in PVD (**Figure 1C, 1D**). This strain enabled us to achieve reliable, unambiguous visualization of PVD and FLP dendrites to investigate their dendritic field sizes, which revealed that dendrites of the FLP neuron cover 23% of the worm’s anterior-posterior longitudinal axis, whereas those of the PVD neuron cover 76% of the worm’s longitudinal axis, with small gaps in coverage between PVD and FLP neurons constituting the final 1% (**Figure 1E**). Our intersectional labeling strategy provides a unique opportunity to elucidate the molecular mechanisms that regulate the boundaries of these dendritic field sizes that collectively span almost the entire length of the worm.

**Figure 1.**
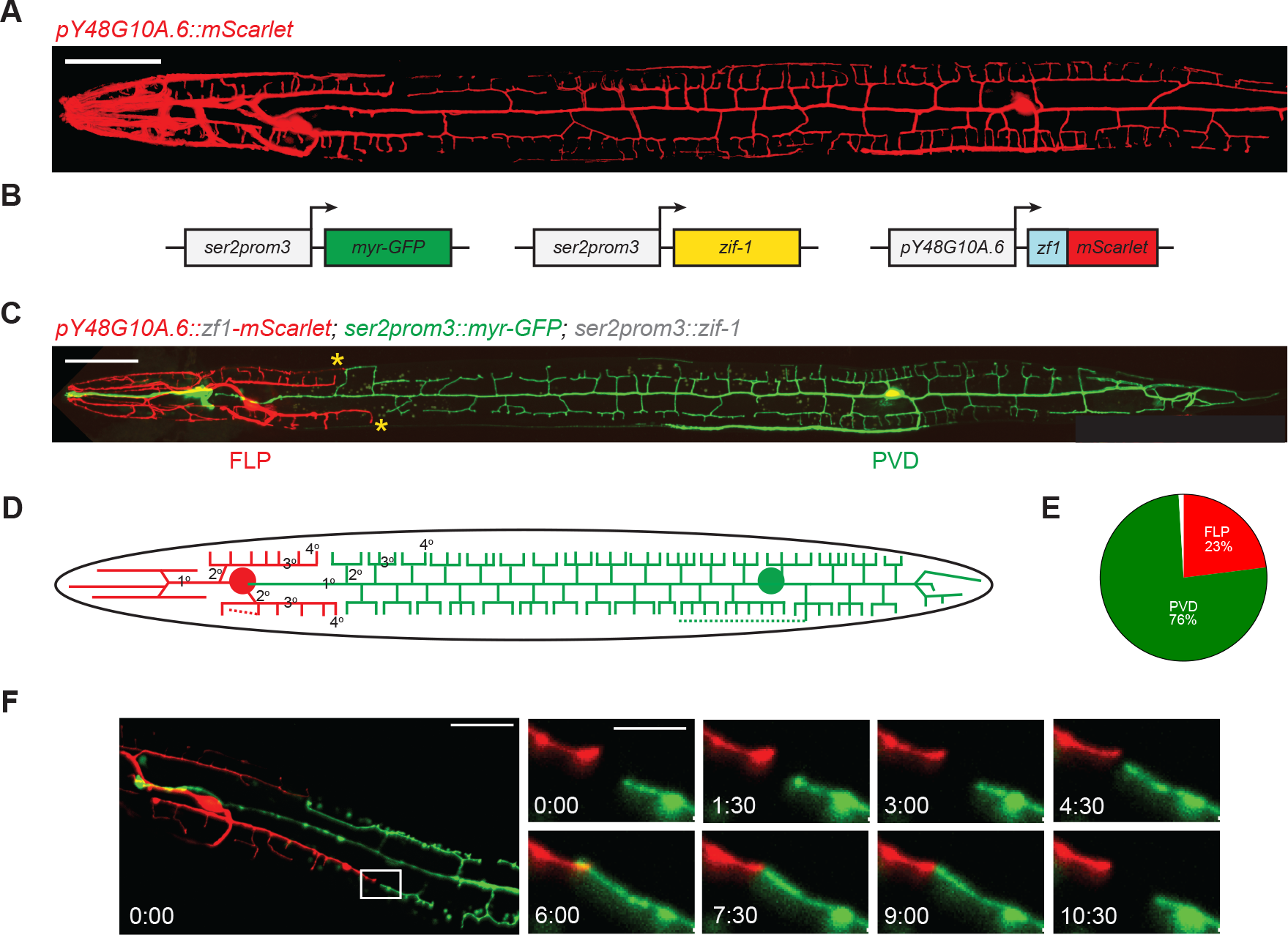
Visualization of the dendritic fields of FLP and PVD mechanosensory neurons. **A)** Visualization of FLP and PVD morphology using transgenic expression of *mScarlet* driven by the FLP and PVD-specific promoter of *Y48G10A.6* (*pY48G10A.6::mScarlet*). Left = Anterior; Top = Dorsal. Scale bar = 50 μm. **B)** Intersectional strategy to achieve cell-specific expression of mScarlet in FLP and myristolated GFPnovo-2 in PVD by co-injection of three transgenes: *ser2prom3::myr-GFP* labels PVD morphology in green; *ser2prom3::zif-1* overexpresses *zif-1* in PVD (ZIF-1 is a SOCS-box adaptor protein that binds to zf1 domains, and targets zf1-containing proteins to an E3 ubiquitin ligase complex for proteasome-mediated degradation); and a *pY48G10A.6::zf1::mScarlet* construct that results in *mScarlet* expression in FLP due to its ZIF-1-mediated degradation in PVD. **C)** Image of an L4 stage worm displaying the morphology of both FLP and PVD neurons, with expression of mScarlet in FLP and GFP in PVD (FLP>mScarlet; PVD>GFP). Yellow asterisks show small gaps in coverage and tiling between FLP and PVD 3° dendrites. The GFP-labeled neuron anterior to the FLP soma is OLL, which is also labeled by *ser2prom3::myr-GFP*.Scale bar = 50 μm. **D)** Schematic of FLP (*red*) and PVD (*green*) morphology. Circle = cell body. Dashed line = axon. **E)** Pie chart depicting percent coverage of the worm’s body in the longitudinal anterior-posterior axis by PVD and FLP dendrites at the L4 stage. Gaps in coverage between PVD and FLP dendrites represent the final 1%. Data represent the mean of n=20 worms. **F)** Frames captured from dynamic imaging movies of growing FLP and PVD 3° dendrites during the L4 stage depict repulsion following contact of neighboring 3° dendrites. White inset indicates the magnified regions displayed on the right, and numbers indicate time in minutes and seconds. Left = Anterior; Top = Dorsal. Scale bars: Left = 50 μm; Right = 5 μm.

### Dendritic tiling by contact-induced retraction between FLP and PVD neurons

These mechanisms underlying dendritic field size regulation are likely related to those that have been documented for tiling of neuronal processes. There have been myriad reports of neurite tiling that arise due to retraction of axons or dendrites that are growing towards one another (Grueber and Sagasti, 2010; Sagasti et al., 2005). However, this is a challenging occurrence to observe. In fact, growing PVD and FLP dendrites have been suggested to not tile by contact-dependent repulsion because laser ablation of either neuron did not result in expanded dendritic territories of the surviving neuron (Yip and Heiman, 2016). To understand if contact-dependent repulsion is present between FLP and PVD dendrites, we sought to image it directly. We first imaged FLP-PVD development to find a developmental timepoint when their dendrites might contact each other (**Figure S1**).

Consistent with a previous report that monitored FLP development, (Androwski et al., 2020), we find that FLP is born embryonically, and by larval stage 1 (L1) a two-pronged, pitchfork shaped primary (1°) has emerged anteriorly from the cell body, as well as a ventral axon. By larval stage 4 (L4), two tertiary (3°) dendrites, one dorsal and one ventral, grow posterior to the FLP soma; these tile with anterior PVD 3° dendrites (**Figure 1D**). Therefore, we anticipated any contact-dependent tiling would occur during L4.

PVD and FLP 3° dendrites grow toward each other in the same linear path because their dendrites are both guided by the ligand SAX-7 on the epidermis that binds to dendritic guidance receptor DMA-1 in PVD and FLP (Dong et al., 2015; Liang et al., 2015; Liu and Shen, 2012; Salzberg et al., 2013). Loss of either *dma-1* or *sax-7* results in severely compromised dendritic growth that eliminates the normally tiled, mosaic pattern of PVD and FLP dendritic fields (**Figure S2A, B**). These results suggest that the precise guidance of PVD and FLP 3° dendrite growth primes them to contact each other (**Figure 1C, S1**).

To determine if FLP and PVD 3° dendrites tile by retracting following contact with each other, we performed *in vivo* dynamic imaging of growing FLP and PVD dendrites. Despite the generally stereotyped development of *C. elegans*, we found contact of PVD and FLP 3° dendrites to occur variably within the L4 stage. Technical limitations restricted our imaging of growing FLP and PVD dendrites during mid-L4 to four hours. Most of our movies showed FLP and PVD dendrites that did not make contact within the four-hour imaging sessions, likely due to the stochasticity of contact during development. However, a small subset of our videos captured points of contact between FLP and PVD 3° dendrites. In these cases, contact-dependent mutual repulsion between FLP and PVD 3° dendrites was observed within a few minutes of contact between FLP and PVD (**Figure 1F**).

### Secreted Wnts CWN-1 and CWN-2 restrict FLP dendritic fields

To identify molecular regulators of dendritic field size confinement, we looked to regulators of PVD dendrite self-avoidance. Sister branches within the PVD arbor avoid each other, and similar molecules might also function to regulate heteroneuronal tiling between adjacent neuronal branches. A previous study discovered that loss-of-function mutants of the Wnt secretory factor MIG-14/Wntless have self-avoidance defects where 10-13% of PVD 3° dendrites overlap (Liao et al., 2018). However, loss of *mig-14* did not result in FLP-PVD tiling defects (**Figure 2A**), since contact-dependent repulsion between FLP and PVD is likely still intact in this case. Rather, FLP dendritic fields exhibited a marked expansion of territory, and the 3° dendrites overgrew further posterior than usual, as quantified by the length of the FLP 3° dendrites that are posterior to the cell body (**Figure 2B**).

**Figure 2.**
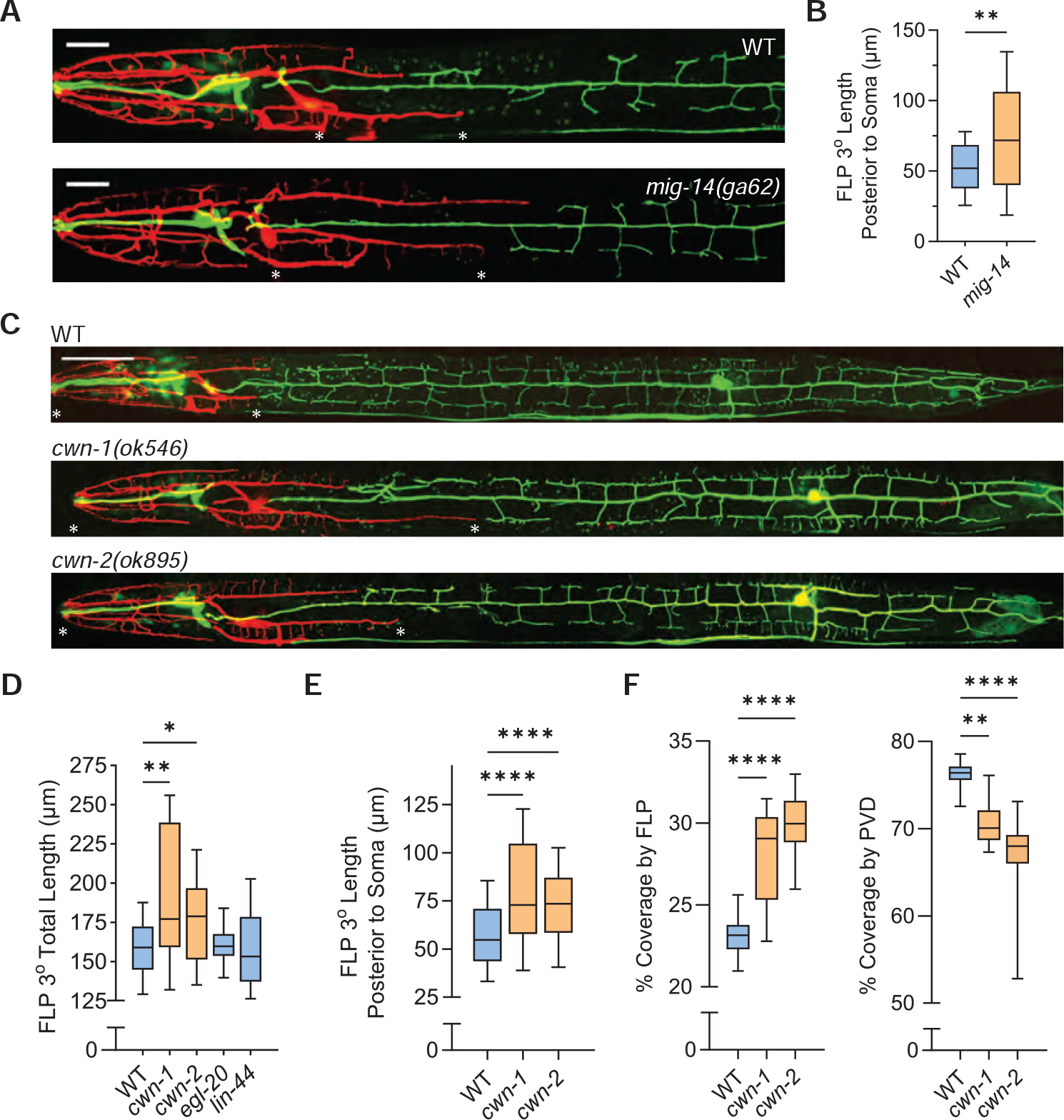
Wnts CWN-1 and CWN-2 regulate relative FLP and PVD dendritic field sizes. **A)** *mig-14*(*ga62*) mutants during the L4 stage exhibit expanded FLP dendritic fields relative to wild-type (WT) worms. Asterisks show length of FLP 3° dendrites posterior to the soma. Scale bar = 20 μm. **B)** Quantification of FLP 3°length posterior to the soma reveals significantly increased FLP 3° dendrite length in *mig-14*(*ga62*) mutants relative to WT. Box, 25%–75% interquartile range; whiskers, 10%–90% interquartile range. ** p < 0.01, unpaired two-tailed *t*-test. WT: n=25, *mig-14*: n=21 worms. **C)** Confocal images of WT, *cwn-1(ok546)*, and *cwn-2(ok895)* mutants during L4. Scale bar = 50 μm. **D)** Loss of Wnts *cwn-1* and *cwn-2*, but not *egl-20* or *lin-44*, results in posterior expansion of FLP dendritic field sizes, set by the total length of FLP 3° dendrites marked by the asterisks in (C). WT: n=21, *cwn-1*: n=17, *cwn-2*: n=28, *egl-20*: n=13, *lin-44*: n=18 worms. **E)** Quantification of FLP 3° length posterior to the soma reveals that this is the major determinant of significantly increased FLP 3° dendrite length and field sizes in *cwn-1(ok546)* and *cwn-2(ok895)* mutants relative to WT worms. WT: n=57, *cwn-1*: n=33, *cwn-2*: n=41 worms. **F)** FLP dendritic fields cover a significantly higher percentage of the worm’s body in *cwn-1(ok546)*, and *cwn-2(ok895)* mutants relative to WT (*Left*), whereas PVD dendritic fields coordinately cover a significantly lower percentage of the worm’s body in *cwn-1(ok546)*, and *cwn-2(ok895)* mutants relative to WT (*Right*). WT: n=20, *cwn-1*: n=18, *cwn-2*: n=14 worms.

MIG-14/Wntless is important for the normal transport of hydrophobic Wnts from the Golgi to plasma membrane. We therefore next tested the hypothesis that secreted Wnts restrict the boundary of FLP dendritic fields. Of the four non-embryonic-lethal Wnts expressed in the *C. elegans* genome (Barstead et al., 2012), *cwn-1* and *cwn-2* loss-of-function mutants, but not *egl-20* or *lin-44*, phenocopied the expanded FLP dendritic arbor phenotype seen in the *mig-14* mutant (**Figure 2C, 2D, 2E**). Because *cwn-2* mutants result in an anterior displacement of the nerve ring, located anterior to the FLP soma (Kennerdell et al., 2009), we also compared FLP somatic positioning of wild-type and *cwn-2* mutants but found no difference (**Figure S2D**). Thus, the significantly longer FLP 3° dendrites seen in the *cwn-2* mutant are not due to an anterior shift in the positioning of the FLP soma and a compensatory attempt to maintain normal dendritic field sizes by overgrowing FLP 3° dendrites posteriorly. Furthermore, both *cwn-1* and *cwn-2* mutants exhibit a significant expansion of FLP coverage at the expense of PVD coverage (**Figure 2F**). These data demonstrate that the Wnts CWN-1 and CWN-2 are required for limiting the growth of FLP dendrites, but are dispensable for the contact-dependent repulsion in tiling because FLP and PVD neurons did not exhibit overlapping or fasciculating dendrites in either of these Wnt mutants.

### CWN-2 puncta are pharyngeal and near the FLP-PVD border

To begin to understand how CWN-1 and CWN-2 may regulate FLP dendritic fields, we aimed to determine where these two Wnts are localized by endogenously tagging both CWN-1 and CWN-2 with GFPnovo2 using CRISPR/Cas9 (Hendi and Mizumoto, 2018). While we successfully knocked-in GFPnovo2 to both Wnts at their endogenous loci, we were unable to detect GFP-tagged CWN-1 (data not shown). At L4, CWN-2/Wnt-GFP puncta were visible at the most anterior portion of the posterior bulb of the pharynx, spreading both anteriorly and posteriorly from this source (**Figure 3A, 3D**). We additionally used a C-terminal mNeonGreen (mNG) tag due to prior reports of its higher intensity (Shaner et al., 2013), and to be consistent with other endogenously tagged Wnts in *C. elegans* (Heppert et al., 2018; Pani and Goldstein, 2018). We saw an equivalent patterning of CWN-2/Wnt throughout the worm using both mNG and GFPnovo2 fluorescent tags (**Figure 3A, 3D**)

**Figure 3.**
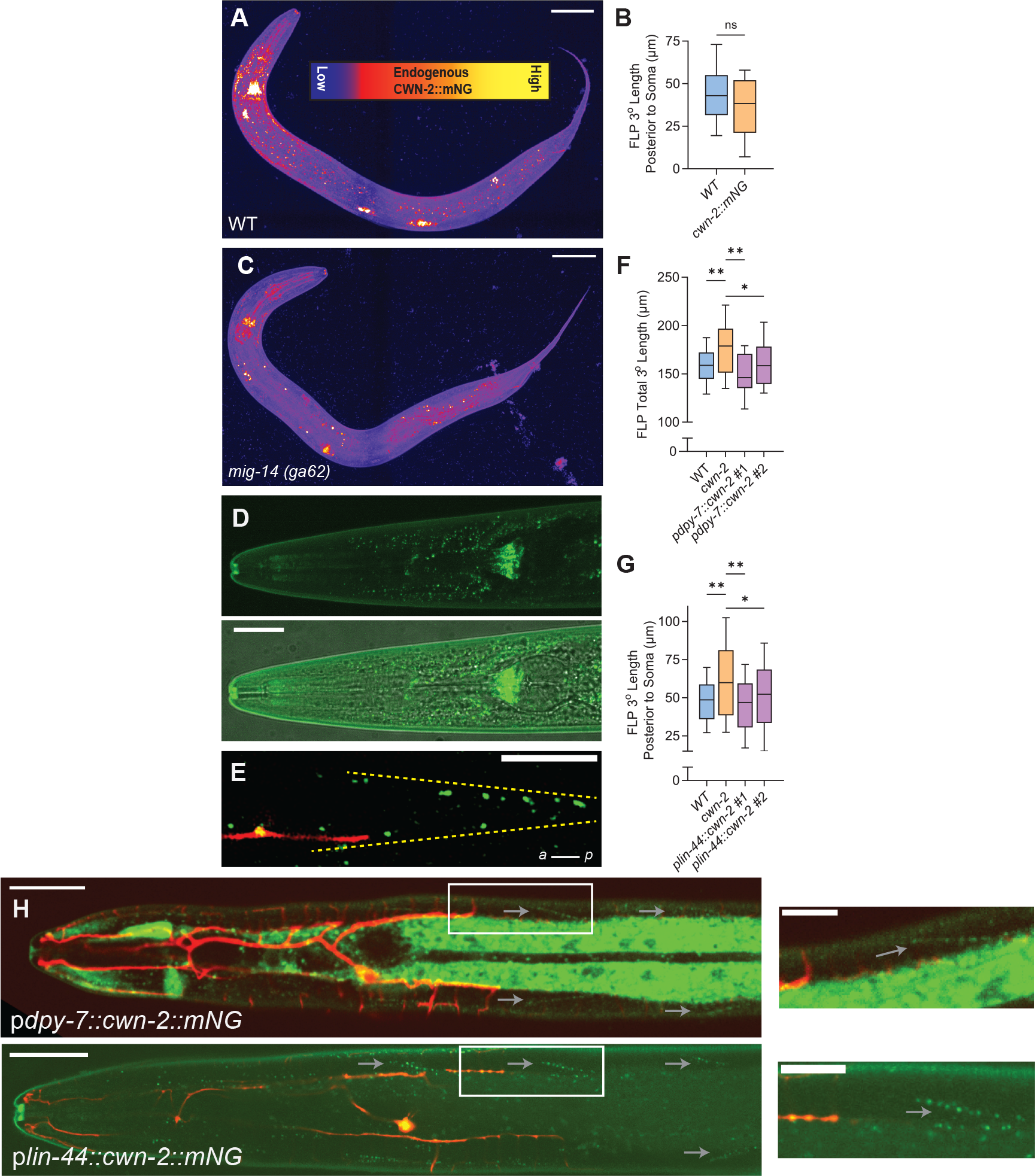
Endogenously tagged Wnt/CWN-2 is in the same plane as growing FLP dendrites. **A)** Representative maximum intensity projection of Wnt/CWN-2::mNeonGreen (mNG) in a late L4 worm pseudo-colored with a Fire lookup table where yellow indicates high intensity (posterior pharyngeal bulb, coelomocytes) and purple indicates low intensity of tagged CWN-2 puncta. Scale bar = 50 μm. **B)** Quanitification of FLP field size in WT and endogenously tagged Wnt/CWN-2::mNG worms demonstrates statistically equivalent FLP dendritic fields and a biologically functional tagged CWN-2 protein. WT: n=19, *cwn-2::mNG*: n=25 worms. **C)** *mig-14(ga62)* mutants display a pronounced loss of punctate CWN-2 throughout the entire body of the worm, as well as a marked reduction in intensity of the CWN-2 at the pharyngeal source relative to WT worms in *(A)*. Scale bar = 50 μm. **D)** Representative maximum intensity projection of Wnt/CWN-2::GFPnovo2 in the head of an L4 worm (*Top*) shows a similar pharyngeal source of CWN-2 to that in Wnt/CWN-2::mNG worms. The overlaid DIC image (*Bottom*) highlights the dominance of CWN-2 puncta in the most anterior portion of the posterior pharyngeal bulb. Scale bar = 25 μm. **E)** Representative image (not maximum intensity projection) of an FLP 3° dendrite (*red*) growing posteriorly in the same plane towards CWN-2::mNG puncta, which are in an elongated arrowhead-shaped pattern posterior to the FLP soma. Dashed yellow lines highlight this pattern. Scale bar = 5 μm, a = anterior, p = posterior. **F)** Rescue of expanded FLP dendritic field sizes following transgenic overexpression of *cwn-2* in hypodermal cells using the *dpy-7* promoter (p*dpy-7::cwn-2)*, from two independent transgenic lines. WT: n=21, *cwn-2*: n=28, p*dpy-7::cwn-2 #1*: n=12, p*dpy-7::cwn-2 #2*: n=21 worms. **G)** Rescue of expanded FLP dendritic field sizes following transgenic overexpression of *cwn-2* in the tail using the *lin-44* promoter (p*lin-44::cwn-2)*, from two independent transgenic lines. WT: n=20, *cwn-2*: n=31, p*lin-44::cwn-2 #1*: n=17, p*lin-44::cwn-2 #2*: n=18 worms. **H)** Representative images (not maximum intensity projections) of L4 worms expressing transgenic p*dpy-7::cwn-2::mNG (Top)* and p*lin-44::cwn-2::mNG (Bottom)*, which display mNG arrowhead patterns (gray arrows) similar to endogenous CWN-2::mNG that are in the same plane as growing FLP 3° dendrites (*red*). Inset on the left is magnified on the right. Scale bars = 25 μm (*Left*), 10 μm (*Right*).

The CWN-2::mNG tag is functional as it did not result in expanded FLP dendritic fields that would indicate a dysfunctional protein (**Figure 3B**). The brightest CWN-2/Wnt dots collectively form a pyramid-shaped pattern just anterior to the most anterior g1 gland cells (**Figure S3A**) (Smit et al., 2008). We speculate that the source of CWN-2/Wnt is likely the pm5 pharyngeal muscle cells given their triangular shape and proximity to the terminal bulb grinder (Altun and Hall, 2009). While this source of CWN-2/Wnt in the posterior pharynx has been previously reported by tagged overexpression constructs of CWN-2/Wnt (Hayashi et al., 2009; Kennerdell et al., 2009), we additionally observed the predominance of CWN-2/Wnt posterior to the terminal bulb of the pharynx, arranged in a series of elongated arrowhead shapes (**Figure 3A, 3E, S3B, S3C**) likely to represent junctions between interlocking pairs of rhomboid-shaped muscles that line the body wall (Mackenzie et al., 1978).

Another prominent localization of CWN-2::mNG is the coelomocytes, which was also not described previously (Hayashi et al., 2009; Kennerdell et al., 2009). In *C. elegans*, coelomocytes are three pairs of macrophage-like cells adjacent to the worm musculature; two are in the head, another two near the midbody, and the final two are in between the tail and the vulva. Indeed, bright expression of *cwn-2::mNG* is visible in all six coelomocytes (**Figure 3A**). In *mig-14 (ga62)* mutants where Wnt is not transported to the plasma membrane for secretion, the intensity of CWN-2::mNG in coelomocytes is indeed significantly reduced, as is that of the puncta in and around the pharyngeal bulb (**Figure 3C**).

The arrowhead localization of CWN-2/Wnt seen in the midbody of the worm is likely to be most relevant to the mechanism by which CWN-2/Wnt and FLP dendrites interact. Intriguingly, these dots of CWN-2/Wnt lie in the direct path of growing FLP posterior 3° dendrites (**Figure 3E**). Based on the loss of function phenotype and the localization of CWN-2/Wnt, we hypothesize that CWN-2/Wnt may act as a molecular “speed bump” for growing FLP dendrites. Thus, fluorescent tagging of *cwn-2* enabled us to visualize the exact endogenous localization of CWN-2.

### CWN-2 is permissive for FLP dendritic field restriction

We next wanted to determine whether CWN-2/Wnt acts as an instructive or permissive cue for growing FLP dendrites. First, we expressed *cwn-2* in the *cwn-2* loss-of-function mutant background under the control of the *dpy-7* promoter (p*dpy-7::cwn-2*). This construct exogenously expresses *cwn-2* in hypodermal cells hyp-5, hyp-6, and hyp-7 that are localized along the body of the worm (Gilleard et al., 1997). Two independent transgenic lines were capable of rescuing the expanded FLP field size seen in *cwn-2* mutants (**Figure 3F**). Furthermore, we additionally cloned and expressed a p*lin-44::cwn-2* transgene in *cwn-2* mutant worms, resulting in CWN-2/Wnt that localizes to epidermal cells in the tail region (Hilliard and Bargmann, 2006; Zheng et al., 2015). Two independent transgenic lines were again capable of rescuing the expanded FLP field size seen in *cwn-2* mutants (**Figure 3G**). Even the mis-expression of CWN-2/Wnt far away from its wild-type pharyngeal source is capable of rescuing FLP dendritic field size back to wild-type lengths. Indeed, mNG-tagged versions of these constructs exhibit a CWN-2::mNG arrowhead pattern in the growing plane of FLP 3° dendrites, similar to that seen in the endogenously tagged strain (**Figure 3H**). Together, these data show that ectopic expression of *cwn-2* from distal sources can recapitulate native CWN-2/Wnt localization patterns to permit the confinement of FLP dendritic fields.

### LIN-17/Frizzled functions in FLP to confine dendritic field size

Because Wnt is a secreted glycoprotein that binds to Frizzled receptors, we next determined whether loss-of-function mutants in *C. elegans* Frizzled receptors might also result in expansions of FLP dendritic territory. Of the three canonical, non-embryonic lethal Frizzled mutants in *C. elegans*, we find that *lin-17* mutants, but not *cfz-2* or *mig-1* mutants, result in FLP dendritic field sizes that were significantly larger than wild-type (**Figure 4A**). These expanded FLP dendritic fields in *lin-17* mutants cover a significantly increased proportion of the worm in the anterior-posterior axis relative to wild-type worms (**Figure 4B, *Left***) (WT: 23% ± 0.4%, *lin-17*: 25.8% ± 0.5%). Expansion of FLP territory in *lin-17* mutants is coordinately accompanied by a significant shrinkage of PVD territory (WT: 76% ± 0.5%, *lin-17*: 68.8 ± 1.5%) (**Figure 4B, *Right***). This finding points to a potential receptor-ligand interaction that together cooperates to maintain the boundaries of FLP neurons.

**Figure 4.**
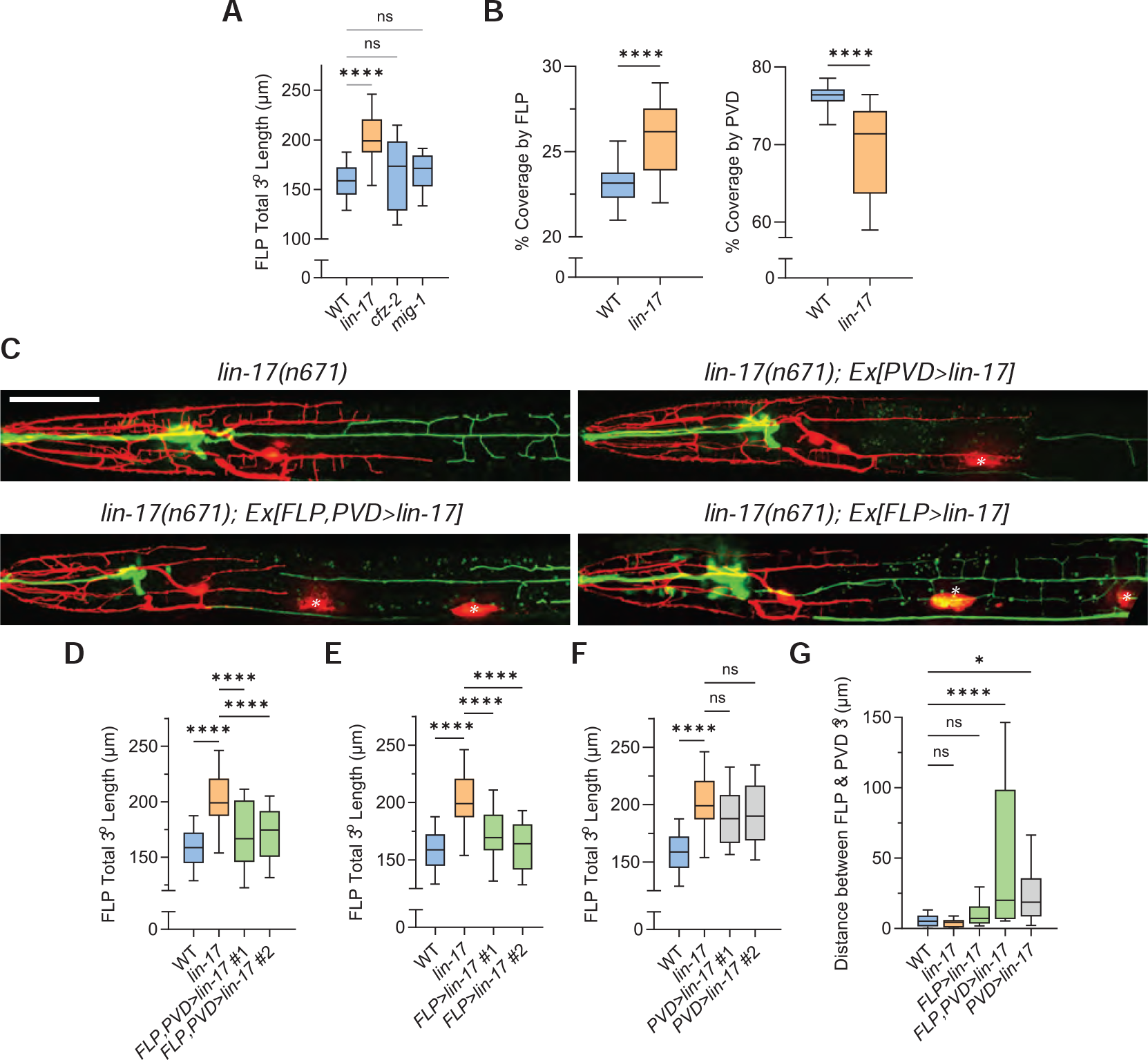
LIN-17/Frizzled functions cell-autonomously in FLP to restrict dendritic field size. **A)** *lin-17(n671)/Frizzled* mutants exhibit significantly expanded FLP dendritic field sizes that phenocopy *cwn-2* mutants, whereas *cfz-2* and *mig-1* mutants do not. WT: n=22, *lin-17*: n=23, *cfz-2*: n=23, *mig-1*: n=27 worms. **B)** FLP 3° dendrites cover a significantly higher percentage of the worm’s body in the anterior-posterior axis in *lin-17* mutants relative to WT worms (*Left*). PVD 3° dendrites coordinately cover a significantly lower percentage of the worm’s body in the anterior-posterior axis in *lin-17* mutants relative to WT worms (*Right*). WT: n=20, *lin-17*: n=25 worms. **** p < 0.0001, unpaired two-tailed *t*-test. **C)** Confocal images of *lin-17(n671)* mutant worms (*Top Left*), worms expressing *lin-17* in both FLP and PVD in the *lin-17* mutant background (*Bottom Left*), worms expressing *lin-17* in only PVD in the *lin-17* mutant background (*Top Right*), and worms expressing *lin-17* in only FLP in the *lin-17* mutant background (*Bottom Right*). White asterisks = p*unc-122::RFP* expression in coelomocytes (co-injection marker). Scale bar = 50 μm. **D-F)** Quanification and analysis of FLP total 3° dendrite length reveal that *lin-17* expression in both PVD and FLP (*D*) or only FLP (*E*), but not in PVD (*F*), rescues FLP dendritic field size in the *lin-17(n671)* mutant background. **G)** Transgenic overexpression of *lin-17* in PVD in the *lin-17(n671)* mutant background (*two right-most boxplots*) induces significant gaps in coverage between FLP and PVD 3° dendrites. WT: n=22, *lin-17*: n=23, *FLP>lin-17*: n=23-26, *FLP,PVD>lin-17*: n=21-24, *PVD>lin-17*: n=27-28 worms. For (*A*), (*D-G*): ns, p > 0.05; * p < 0.05, **** p < 0.0001 one-way ANOVA and Tukey’s multiple comparisons test.

To determine whether LIN-17/Frizzled might interact with CWN-2/Wnt to restrict FLP growth, we first generated an endogenous, C-terminal GFPnovo2 knockin of *lin-17* to visualize its expression pattern (**Figure S4A**). This LIN-17-GFPnovo2 strain exhibited strong expression near the vulva and tail, consistent with previous studies (Klassen and Shen, 2007; Kurshan et al., 2018). Despite dim but visible expression in both anterior and posterior pharyngeal bulbs, limited expression was seen in the FLP cell body or its 3° dendrites that determine its posterior boundary.

Low receptor expression may still be sufficient for function in FLP; therefore, we turned to a transgenic overexpression approach to determine *lin-17*’s site of action. First, we expressed *lin-17* in both PVD and FLP neurons (p*Y48G10A.6::lin-17*) in a *lin-17Δ* mutant background (**Figure 4C, 4D**). FLP and PVD expression of *lin-17* was sufficient to rescue the expanded FLP arbor in a *lin-17Δ* mutant. Rescue of the *lin-17* mutant phenotype was also achieved by expression of *lin-*17 only in FLP (**Figure 4E**), achieved by linking a zf-1 tag to LIN-17 and ZIF-1 expression in PVD (*ser2prom3::zif-1*, **Figure 1B**). However, in the *lin-17* mutant background, exogenous expression of *lin-17* only in PVD did not result in the rescue of FLP dendritic field size; rather, its overexpression results in significant gaps in dendritic coverage between FLP and PVD 3° dendrites at the L4 stage (**Figure 4F**). These gaps between neighboring FLP and PVD 3° dendrites induced by exogenous *lin-17* expression in PVD are additionally accompanied by pronounced defects in PVD morphology, including significant shortening of the PVD 1° dendrite and decreased density of PVD 2° dendrites that are not seen in the *lin-17Δ* mutant alone (**Figure S4B-D**).Taken together, these results demonstrate that *lin-17* likely functions in FLP to restrict its dendritic 3° growth, and that *lin-17* is either not expressed in PVD or is less sensitive to CWN-2/Wnt in PVD relative to FLP.

### CWN-2 slows growth of FLP dendrites to limit dendritic field size

Having identified a set of molecular components of FLP dendritic field size, we were interested in understanding how exactly CWN-2/Wnt might restrict FLP dendritic fields. We hypothesized that CWN-2/Wnt acts as a repulsive cue for FLP dendrites, causing slowed growth and/or retraction upon LIN-17/Frizzled activation. To test this hypothesis, we performed dynamic imaging of CWN-2::mNG worms *in vivo* during growth of FLP at L4. Growth of both FLP 3° and 4° dendrites was apparent at this stage, especially when CWN-2/Wnt puncta were not in the vicinity (**Figure 5A, 5C**). However, we noticed a striking slowing of FLP dendritic growth when dendrites contacted CWN-2/Wnt. FLP would contact CWN-2/Wnt puncta and either completely retract, or retract then resample the puncta over a period of a few minutes. In other cases, FLP would contact CWN-2/Wnt, and remain bound for several minutes before the resumption of forward growth (**Figure 5B, 5D**). Consistent with these observations, the net growth and speed of FLP dendrites is slowed upon contact with CWN-2/Wnt, as shown by both distance traveled by growing FLP dendrites (**Figure 5E**), the cumulative probability distributions of growth/retraction events over each 30 second frame (**Figure 5F**), and average speed of FLP dendrites with and without contact of CWN-2::mNG (**Figure 5G**). Taken together, our data indicate that CWN-2/Wnt acts as an inhibitor of forward FLP dendrite growth by inducing retraction and/or pausing of growth.

**Figure 5.**
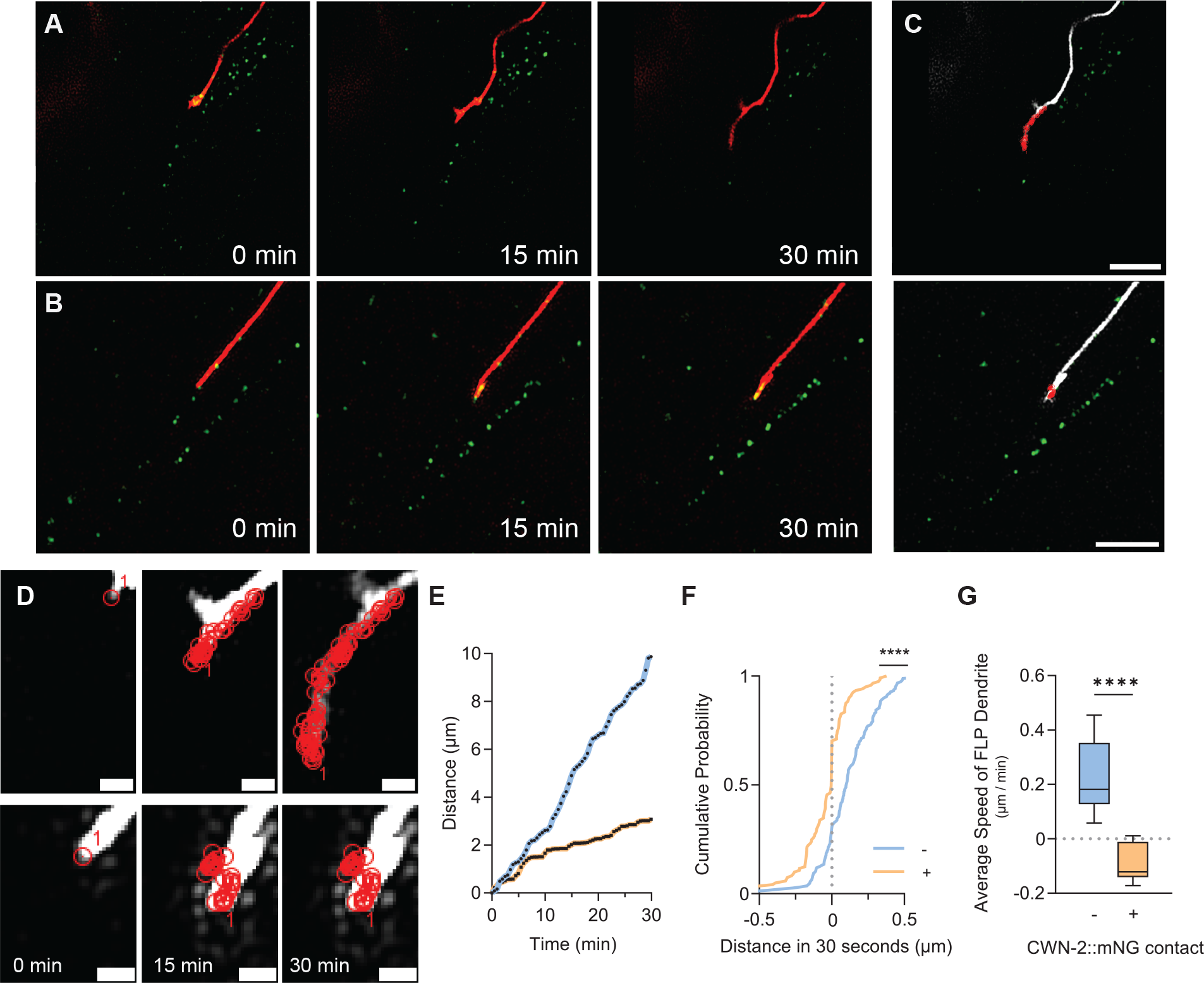
Inhibition of FLP dendrite growth upon CWN-2 contact. **A)** Frames taken from representative dynamic imaging movies of a growing FLP 3° dendrite (*red*) *in vivo* that encounters little to no contact of CWN-2::mNG puncta at the tip of the dendritic growth cone over a thirty minute period. **B)** Frames taken from representative dynamic imaging movies of a developing FLP 3° dendrite *in vivo* that encounters two distinct CWN-2::mNG puncta along its growing path over a thirty minute period. **C)** Small red circles indicate the position of the FLP growth cone at each frame (taken every thirty seconds) for the movie in (*A) (Top*) and in (*B) (Bottom*). The FLP dendrite is pseudocolored white for clarity. Scale bar = 4 μm. **D)** Zoomed-in data in from (*C) Top* and *Bottom* shown with more timepoints for clarity. Red circles indicate the tip of the FLP dendritic growth cone at each timepoint (60 total for a 30min movie). “1” marks the approximate location of the tip of the FLP dendritic growth cone at the indicated timepoint on the bottom left. Scale bar = 1 μm. **E)** Plot of distance (in μm) traveled by the FLP growth cone in (*A, C blue dotted line)* and (*B, D orange dotted line)* relative to the first frame over a thirty minute dynamic imaging session reveals markedly reduced growth in FLP dendrites that contact CWN-2::mNG puncta during their growth trajectory. **F)** Cumulative probability distributions of growth (> 0 μm) or retraction (< 0 μm) over a 30 second interval for FLP dendrites that contact CWN-2::mNG puncta (orange line, “+”) and FLP dendrites that do not contact CWN-2::mNG puncta (blue line, “-”). **** p < 0.0001, Kolmogorov-Smirnov test. “-”: n=192, “+”: n=121 dendritic growth events. **G)** Mean speed of growing FLP dendrites (μm/min, n=21) is significantly reduced upon CWN-2::mNG contact (+) relative to those that do not contact CWN-2::mNG (-). **** p < 0.0001, two-tailed *t*-test.

## Discussion

Here we show that contact-induced retraction does indeed contribute to dendritic tiling between FLP and PVD mechanosensory neurons. However, we find this mechanism does not act alone to restrict dendritic arbors from overgrowing into their neighbors’ territory. Anatomical barrier or molecular cues have been proposed to function in prevent tiling errors (Yip and Heiman, 2016). We have identified one extracellular cue, CWN-1/CWN-2 and LIN-17 signaling, that acts in conjunction with dendro-dendritic repulsion to restrict dendritic field sizes by slowing growth of FLP dendrites through LIN-17/Frizzled (**Figure 6**).

**Figure 6.**
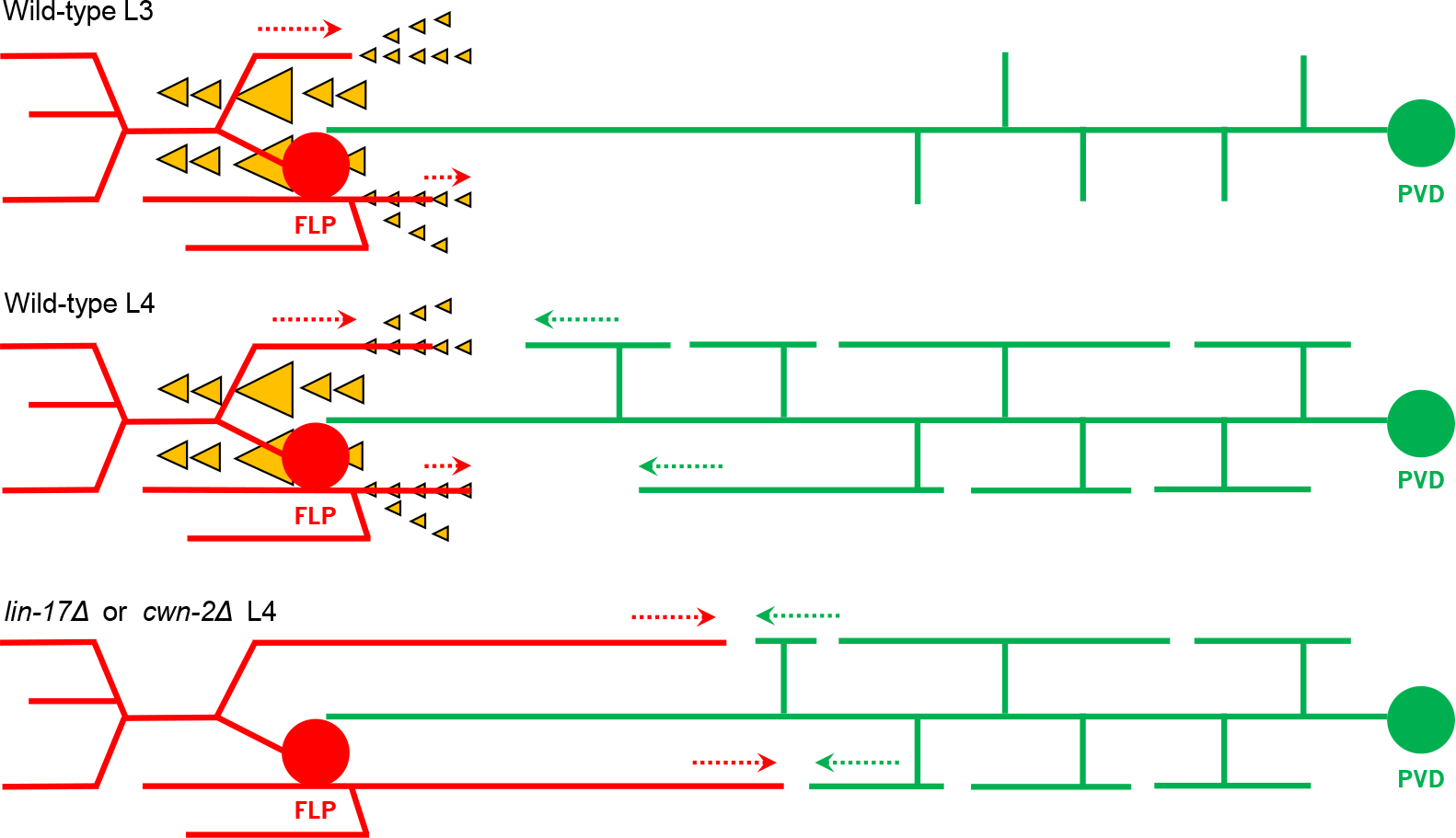
FLP dendritic field restriction by CWN-2/Wnt, LIN-17/Frizzled, and contact-dependent repulsion with PVD dendrites. Prior to L4, FLP 3° dendrites grow posteriorly, but are slowed down upon contact with nearby CWN-2/Wnt puncta due to LIN-17/Frizzled-specific function in FLP. By L4, PVD 3° dendrites that are distal from the PVD soma have finally grown anterior enough, resulting in contact-dependent repulsion between FLP and PVD 3° dendrites that grow linearly along SAX-7 stripes present on the epidermis. With the loss of *cwn-2/Wnt* (*cwn-2Δ*) or *lin-17/Frizzled* (*lin-17Δ*), inhibition of growth is reduced, which enables FLP 3° dendrites to grow further posterior than usual. This unrestricted growth of FLP dendrites results in an expansion of FLP dendritic fields and a reduction of PVD dendritic fields. Yellow triangles indicate CWN-2/Wnt puncta and dashed arrows indicate direction of dendrite growth.

In order to gain insight into possible mechanisms for how Wnts restrict FLP dendritic field size, we endogenously tagged CWN-2/Wnt with two fluorophores. The pattern of CWN-2/Wnt in the present study only partially overlap with previously published reports that utilized tagged overexpression constructs. A *cwn-2::SL2::GFP* bicistronic expression vector revealed expression in the pharynx, intestine, muscles in the anterior body wall, vulva and SMD head neuron (Kennerdell et al., 2009). Another study investigating CWN-2-dependent mechanisms of neurite pruning of the head interneuron AIM found that a 0.8 kb putative promoter region upstream of the *cwn-2* initiation codon fused to GFP resulted in detectable GFP expression in several pharyngeal neurons and in the intestine (Hayashi et al., 2009).

However, the direct visualization of endogenously tagged CWN-2/Wnt in the present study revealed that there is an additional abundance of CWN-2/Wnt protein in the posterior pharynx that spreads to the border of FLP and PVD dendritic arbors tens of microns away. Near this border, CWN-2/Wnt is visible as an extended arrowhead pattern, likely co-inhabiting the space between muscle cells and skin cells that FLP and PVD dendrites also traverse during their growth trajectory. However, CWN-2/Wnt is not confined to the space surrounding the pharynx; it is also visible in other regions of the worm as far away as the posterior third of the worm such as the posterior coelomocytes, forming a long-range pattern that is consistent with other studies that have endogenously tagged other Wnts in *C. elegans* (Heppert et al., 2018; Pani and Goldstein, 2018).

Interestingly, the exact source of CWN-2/Wnt is dispensable for setting the boundaries of FLP dendrites. Overexpression of *cwn-2* elsewhere in the *cwn-2* mutant background, even as far away as the tail, can still rescue FLP field sizes back to wild-type levels. This suggests that posteriorly-expressed CWN-2/Wnt is able to reach the head in a presently unidentified mechanism. The permissiveness of CWN-2/Wnt here is consistent with previous studies of CWN-2/Wnt in nerve ring placement, for example (Kennerdell et al., 2009).

In developing neurites, Wnts have been found to act as chemoattractants in some contexts, and chemorepellents in others (Kolodkin and Tessier-Lavigne, 2011). Our dynamic imaging experiments here revealed that contact with CWN-2/Wnt by FLP dendrites results in retraction and slowed growth. This finding is in line with a study showing that the *Drosophila* homolog of *cwn-2*, Wnt-5, and the receptor tyrosine kinase Derailed/Ryk mediate axonal repulsion away from the posterior commissure (Yoshikawa et al., 2003). Wnts also repel descending motor corticospinal axons in mice through Ryk (Liu et al., 2005). However, CWN-2/Wnt has also been shown to be an attractant for processes of the GABAergic head neurons RMED and RMEV, acting through MIG-1 and CFZ-2 Frizzled receptors (Song et al., 2010), which we found to be dispensable for FLP field size restriction. Different head neurons likely utilize distinct Frizzled receptors to differentially respond to CWN-2/Wnt.

During development, the processes of within-class sensory neurons segregate to occupy distinct territories to completely cover receptive fields. This study yields additional insight into the molecular and developmental mechanisms that regulate sensory receptive fields in *C. elegans* that are covered by the harsh touch-responsive FLP and PVD multi-dendritic neurons. In addition to contact-induced retraction, extracellular cues such as CWN-2/Wnt and cell-specific function of receptors such as LIN-17/Frizzled contribute to defining the boundary between neighboring sensory neurons, perhaps acting as a failsafe to ensure that tiling remains complete and non-redundant. How these secreted cues such as CWN-2/Wnt interact with cell-autonomous adhesion molecules and intrinsic growth receptors will remain a topic for future investigation.

## Materials and Methods

### C. elegans Methods

*C. elegans* strains (**Supplementary Table 2**) were cultured on nematode growth medium (NGM) plates seeded OP50 *Escherichia coli* (Brenner, 1974). N2 Bristol was used as our wild-type strain. *C. elegans* transgenic strains were generated via gonadal microinjection of adult worms of the designed construct at a concentration of 1-5 ng/μL, in addition to a co-injection marker at 30-40 ng/μL for podr-1::GFP and punc-122::RFP, or 2-5 ng/μL for pY48G10A.6::mScarlet. Additional empty vector (pSK+) was co-injected in cases where the aforementioned plasmids to be injected totaled less than 40 ng/μL. Worms for imaging were grown at 20ºC and imaged at room temperature, at the L4 larval stage with the exception of developmental experiments when imaging occurred earlier as noted.

### Constructs and Cloning

All plasmids used in this study for expression in *C. elegans* were generated using pSM delta as a backbone, commercially synthesized oligonucleotides, and isothermal assembly (Gibson et al., 2009). Constructs were validated by restriction enzyme digestion and sequencing. Promoter sequences for fusion constructs in Figure 1 are listed in **Supplementary Table 1**.

### Genome editing using CRISPR/Cas9

Endogenous GFPnovo2 and mNeonGreen insertions were generated by microinjection of CRISPR–Cas9 protein complexes into the gonad of adult worms. CRISPR–Cas9 genome editing was done with injections of (in μM): 1.525 Cas9 protein, 4.5 tracrRNA (IDT), and 4.5 crRNAs (IDT). Both tracrRNA and crRNAs were resuspended in duplex buffer (IDT). DNA repair templates were generated using PCR and injected at greater than 50 ng/μL after melting (Ghanta and Mello, 2020). A crRNA against *dpy-10(cn64)* (crRNA sequence: *gctaccataggcaccacgag*; Repair template: *cacttgaacttcaatacggcaagatgagaatgactggaaaccgtaccgcATgCggtgcctatggtagcggagcttcacatggcttcagaccaacagcct*) was co-injected to generate a dominant roller phenotype as a marker for ‘jackpot broods’ according to published methods (Arribere et al., 2014; Paix et al., 2015). F2 worms on plates with the greatest percentage of rollers were singled onto individual plates, lysed, and successful knockin of the fluorophore was confirmed using PCR followed by sequencing. Worms with the desired genome edit were then outcrossed prior to use for experimentation.

### Confocal Microscopy Imaging/Analysis

Static confocal images of FLP and PVD dendrite morphology, and of fluorescently tagged fusion proteins were taken at room temperature in live worms. L4 stage hermaphrodites were anesthetized using in 10 mM levamisole in M9 buffer on 3% agarose pads. Worms were imaged on a 3i Marianas spinning disk system including an inverted Zeiss Axio Observer Z1 microscope, a Prime 95B Scientific CMOS camera (Photometrics), a Yokogawa CSU-W1 spinning-disk, and a C-Apochromat 40× 0.9 NA water-immersion objective using 3i Slidebook (v6) software. Image settings including laser power and exposure time were identical for all genotypes within each experiment.

Quantification of maximum intensity projections (MIPs) was performed using ImageJ/Fiji (National Institutes of Health) (Schindelin et al., 2012). These MIPs were occasionally rotated and straightened for improved visualization, and brightness was modified to best highlight FLP morphology for quantification of FLP dendritic field sizes. For particular experiments that required comparison of pixel intensity between strains, brightness was adjusted to identical values between all genotypes within the experiment. A fire look-up table was utilized for visualization of CWN2::mNeonGreen localization in Figure 3 with an accompanying heat map generated by E. Nichols.

For calculations of FLP dendritic field size, the distance between the most distal point of the FLP 3° dendrite to the tip of the mouth was measured using a segmented line in Fiji. Each worm thus generates two values, one distal and one ventral, and both were included in our analyses. For calculations of FLP 3° length posterior to the soma, the distance between the most distal point of the FLP 3o dendrite to the most posterior position of the soma was quantified using a segmented line in Fiji.

### Dynamic Imaging

For dynamic imaging experiments in Figure 1 and Figure 5, early to mid L4 stage worms were immobilized using 2.5 to 5 mM levamisole in M9 on 5 to 6% circular agarose pads (1.4 mm diameter) that fit into the center of a 35mm Mattek dish. Movies were recorded for up to four hours with 90 second intervals between frames (Figure 1), or one to two hours with 30 second intervals between frames (Figure 5).

Dynamic imaging of contact-induced retraction (Figure 1) was performed on an inverted Zeiss Axio Observer Z1 microscope and 3i software as described in the previous section. Dynamic imaging of FLP dendritic contact with CWN-2::mNeonGreen (Figure 5) was performed on a single plane (no Z-stacks), with 30 second intervals between frames, using a 63X C-Plan Apochromat 1.42NA objective on a Zeiss LSM980 Airyscan 2 system with 405- and 488-nm solid state lasers. To account for incomplete paralysis with levamisole, we utilized a StackRegJ plugin (Stowers Institute) in Fij/ImageJ, as well as MTrackJ (Meijering et al., 2012) to trace the trajectories and measure the lengths of growing FLP dendrites (Figure 5).

### Statistics & Reproducibility

Data are displayed as boxplots with boxes representing the median and interquartile range (25^th^ to 75%th percentile), and whiskers displaying the 10^th^ and 90^th^ percentiles. Graphs showing dendritic field size or dendrite length include measurements from both FLP dorsal and ventral 3° dendrites. Therefore, in most cases, measurements from a single worm will yield two values and both are included in our analyses and plots as two distinct values.

Statistical analyses were performed with two-tailed *t*-tests (for statistically significant differences between two groups) or a one-way ANOVA with Dunnett’s multiple comparisons test (for statistically significant differences between three or more groups). Samples sizes are noted in the legends for each figure. For all graphs, ****, p < 0.0001; ***, p < 0.001; **, p < 0.01; *, p < 0.05; ns (not significant), p > 0.05. All statistical analyses and graphs were generated in Prism 9.0 (GraphPad Software, LLC).

## Acknowledgements

We thank Drs. T. Stearns and L. Luo for conceptual advice and discussions, and all members of the Shen laboratory for helpful discussions and input on the manuscript. We thank R. Sze for assistance with *lin-17* rescue experiments, V. Paat for excellent technical support, and D. Kawano, E. Nichols, Drs. H. Deng, N. McDonald, and R. Shi, for graphics and specific input on *in vivo* dynamic imaging, analysis. This research was supported by the Howard Hughes Medical Institute (to K.S.), NIH R01 (to K.S., T.S., NS082208), and NIH Postdoctoral Individual National Research Service Award (to C.P.T., F32NS120933).

## Author Contributions

C.P.T., and K.S. designed research; C.P.T. performed research; C.P.T. contributed new reagents/analytic tools; C.P.T analyzed data; C.P.T. and K.S. wrote the paper.

