## Supplemental Material for "Wnt signaling and contact-mediated repulsion shape sensory dendritic fields"

**This PDF file includes:**

Figures S1 to S4
Supplementary Tables 1 to 2

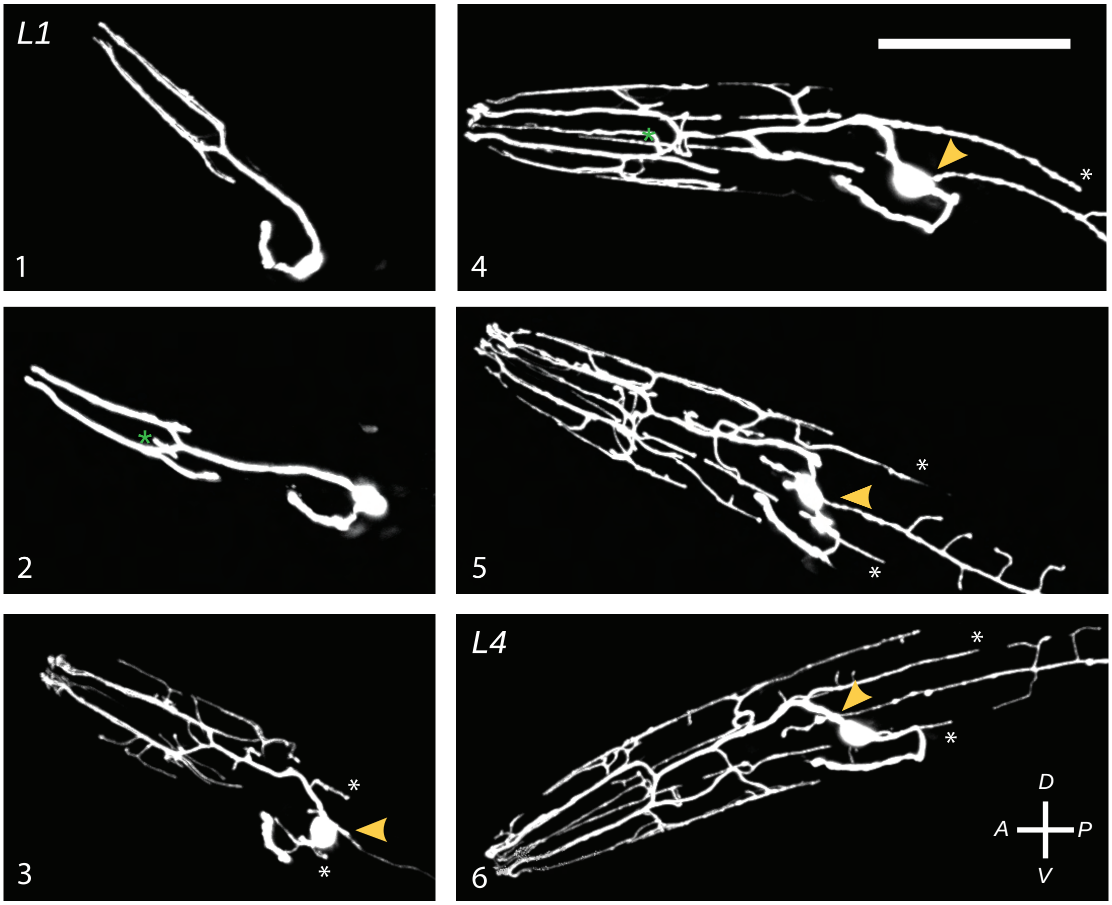

**Supplementary Figure S1. Development of FLP mechanosensory neurons.**

In *C. elegans*, there are two FLP neurons, one on the left and another on the right side. FLP from one side is shown here, pseudocolored white from a p*Y48G10A.6::mScarlet* transgenic strain that labels the morphology of both FLP and PVD neurons. At larval stage 1 (L1), FLP neurons consist of a cell body located near the posterior bulb of the pharynx and a pitchfork-shaped primary (1º) dendrite that extends anterior to the soma, with its ciliated endings terminating near the lateral lips of of the worm's mouth. The FLP axon extends ventrally from the soma. A third dendritic "prong" of the pitchfork then extends in between the two older dorsal and ventral prongs, most visible in panels #2 and #4 (green asterisks). Secondary (2º) dendrites later grow out orthogonal to the base 1º pitchfork structure, followed by tertiary (3º) dendrites that extend orthogonally to the 2º ones; the most posterior of these FLP 3º dendrites (white asterisks) are the ones that most prominently tile with anterior PVD 3º dendrites. Contact-dependent repulsion between FLP and PVD 3º dendrites occurs during L4, as does growth of FLP quaternary (4º) dendrites.

Yellow arrowheads indicate the PVD 1º dendrite growing past the FLP soma. Numbers in bottom left of each panel represent the approxiate age of the animal shown in each panel: 1 is youngest, and 6 is oldest. Scale bar = 50 µm. *A* = anterior, *P* = posterior, *D* = dorsal, *V* = ventral.

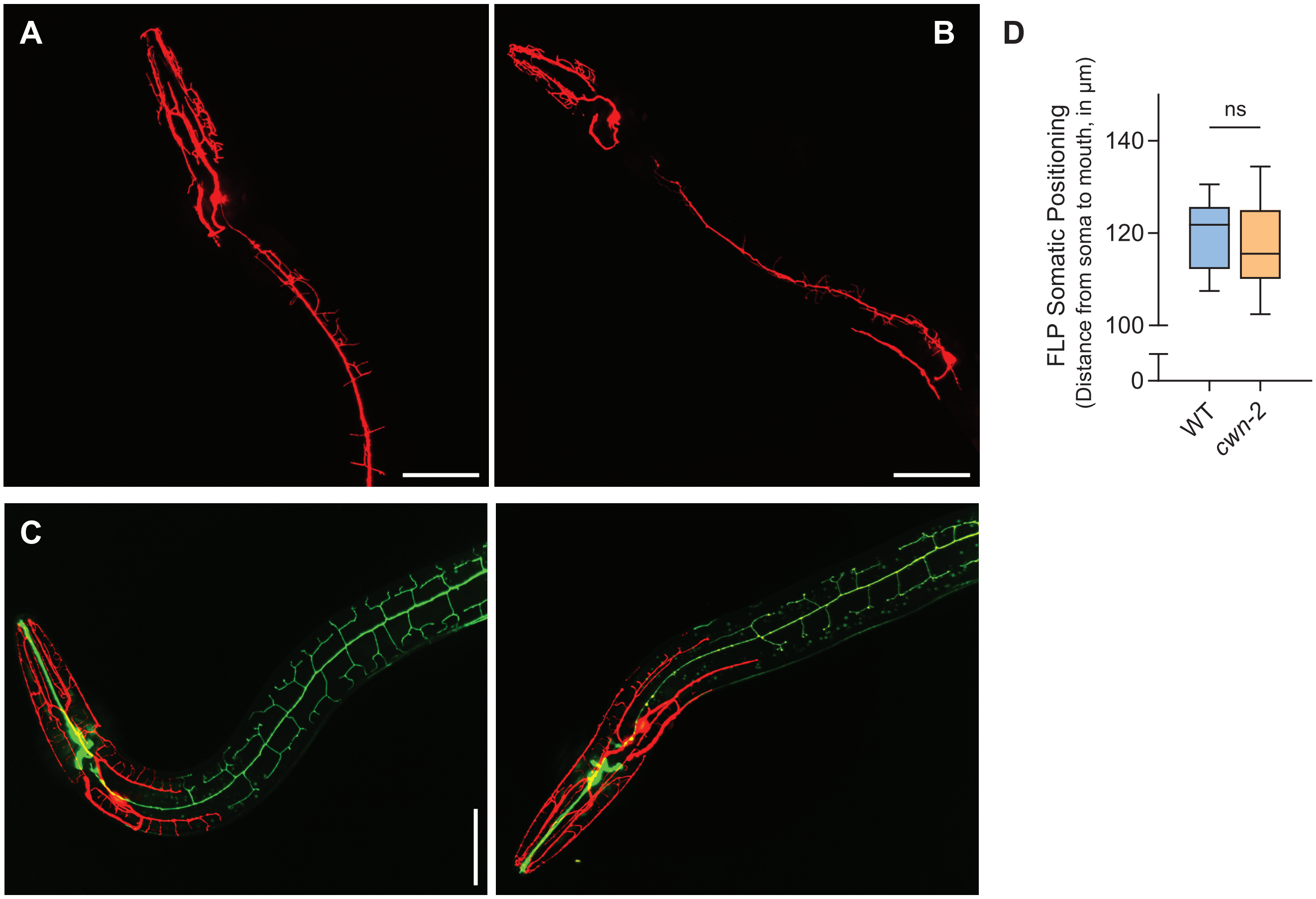

**Supplementary Figure S2. *dma-1* and *sax-7* mutants display severely undergrown FLP / PVD dendrites, whereas *Y48G10A.6* knockout worms do not.**

**A)** *dma-1 (tm5159); pY48G10A.6::mScarlet* mutant worms exhibit significant deficits in FLP and PVD dendrite growth (*Liu and Shen, 2011*), and thus disrupted incomplete coverage.

**B)** Severe deficits in FLP and PVD dendritic growth in *sax-7 (nj48); pY48G10A.6::mScarlet* mutants also result in dendritic morphology and field size abnormalities.

**C)** Deletion of the entire coding region of the *Y48G10A.6* using CRISPR/Cas9 results in normal dendritic tiling between PVD (*green*) and FLP (*red*). Dendritic morphology of *Y48G10A.6* mutants also appears indistinguishable from that of wild-type worms. Two representative *Y48G10A.6* knockout worms are shown. Note: *ser2prom3::GFP* labels the head neuron OLL in addition to PVD.

**D)** No statistically significant difference in FLP somatic positioning between WT and *cwn-2* mutants (p=0.55, two-tailed *t*-test, WT: n=25, *cwn-2*: n=31)

Scale bars = 50 µm.

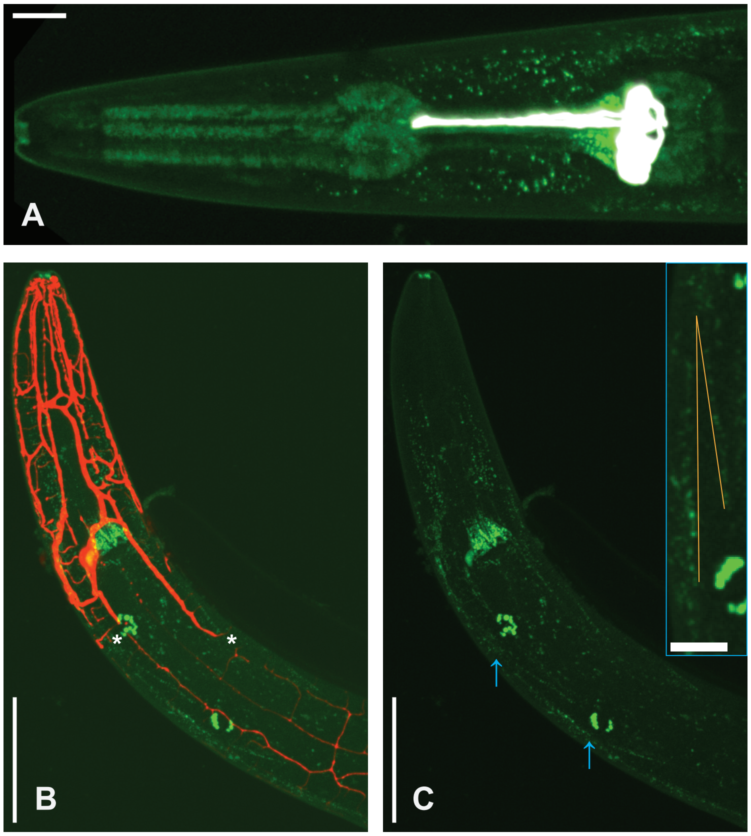

**Supplementary Figure S3. Localization of CWN-2/Wnt *in vivo*.**

**A)** Representative image showing the incomplete colocalization of CWN-2::GFPnovo2 puncta with p*phat-5::mScarlet* expression (pseudocolored white for clarity) in anterior g1 gland cells, g1AR and g1AL (*Smit et al., 2008*). The CWN-2 source is anterior to g1AR and g1AL, likely pm5 pharyngeal muscle cells. Scale bar = 10 µm.

**B)** Representative image displaying CWN-2:mNG puncta near the FLP-PVD border, which is shown by the white asterisks. Both FLP and PVD are visualized with mScarlet in this case for unambiguous CWN-2 visualization. Scale bar = 50 µm.

**C)** CWN-2::mNG puncta that are posterior to the pharynx are arranged in an elongated arrowhead pattern likely representing muscle cell junctions. Blue arrows mark boundaries of the inset, highlighting the arrangement of CWN-2 in an "arrowhead' (yellow lines) with an 8.8º acute angle in this example. Scale bars = 50µm (*Left)*, and 10µm (*Right Inset*).

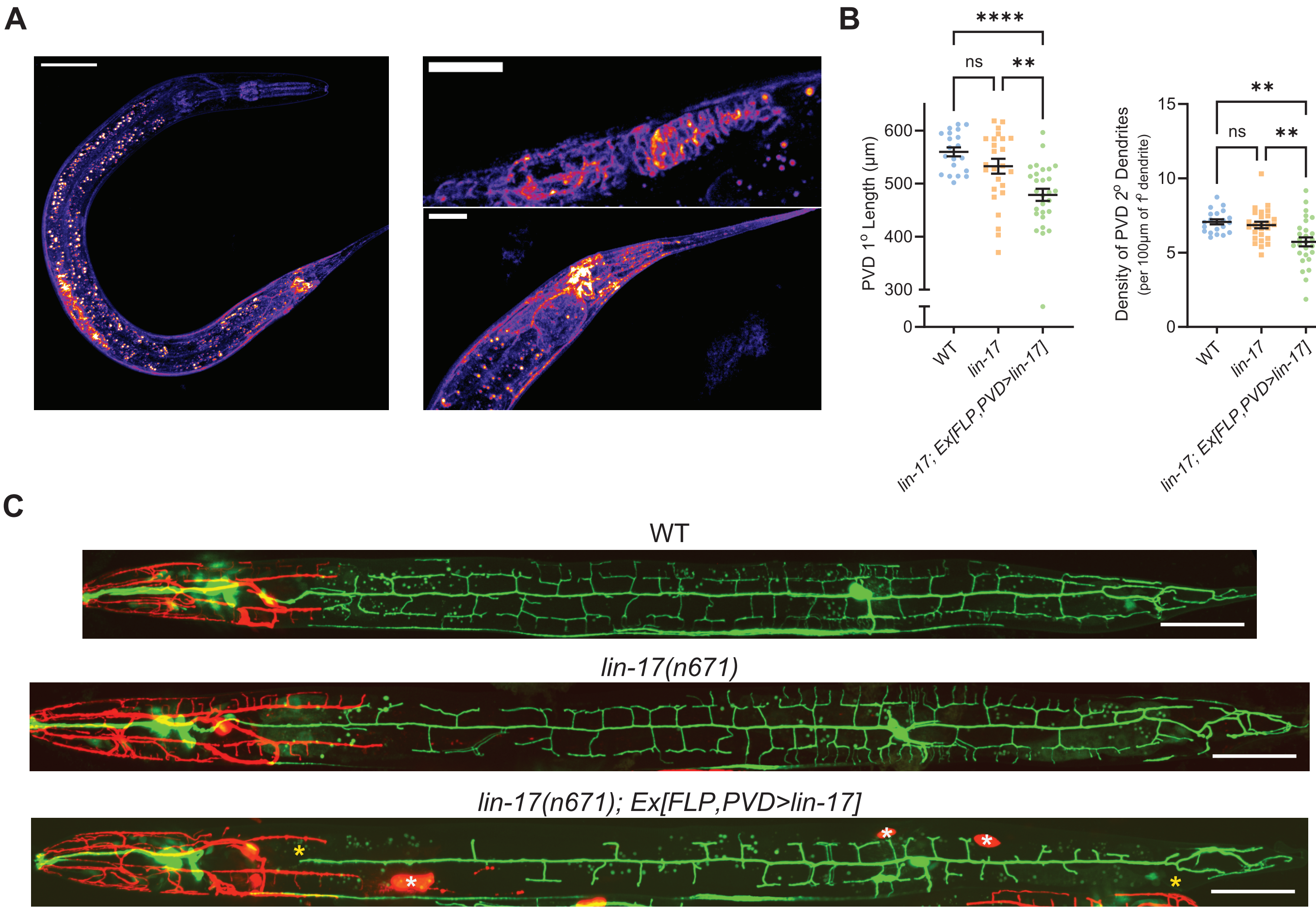

**Supplementary Figure S4. Expression of *lin-17* in PVD disrupts dendritic morphology.**

**A)** Endogenous GFPnovo2 tag of LIN-17 shows high *lin-17::GFPnovo2* expression in the vulva (*Top Right*) and tail (*Bottom Right*). Images on the left and right are from different worms. Scale bars = 50 µm (*Left*), and 20 µm (*Right*).

**B)** Expression of *lin-17* in PVD results in the significantly reduced length of the PVD 1° dendrite (*Left*), and significantly decreased density of PVD 2° dendrites (*Right*). These defects in PVD morphology are not seen in the *lin-17(n671)* mutant.

**C)** Representative images of data shown in (B). Red ovals in the *lin-17(n671); Ex[FLP,PVD>lin-17]* image show coelomocytes expressing the p*unc-122::RFP* co-injection marker (*white asterisks*) used to identify worms with expression of the *Ex[FLP,PVD>lin-17]* transgene. Yellow asterisks highlight the decreased length of the PVD 1° dendrite resulting from the expression of *lin-17* in PVD. Scale bars = 50 µm.

| Promoter | Length (bp) | Fluorophore | log FC | 1% | 2% | padj |
|---|---|---|---|---|---|---|
| FLP | | | | | | |
| *flp-5* | 5,419 | mScarlet | 2.32 | 0.974 | 0.316 | 1.76E-59 |
| *D2096.9* | 4,832 | mScarlet | 1.65 | 0.782 | 0.058 | 1.86E-168 |
| *T10G3.8* | 916 | mScarlet | 1.26 | 0.744 | 0.029 | 6.53E-292 |
| *nlp-66* | 2,224 | mScarlet | 1.06 | 0.41 | 0.008 | 0 |
| *K09E2.1* | 245 | mScarlet | 0.767 | 0.538 | 0.005 | 0 |
| *R05H11.2* | 685 | mScarlet | 0.763 | 0.385 | 0.017 | 1.99E-129 |
| *gst-5* | 1,148 | mScarlet | 0.691 | 0.577 | 0.148 | 9.84E-23 |
| *F46B6.2* | 3,159 | mScarlet | 0.57 | 0.449 | 0.018 | 2.43E-171 |
| *Y71H2AR.2* | 1,242 | mScarlet | 0.563 | 0.423 | 0 | 0.00E+00 |
| *valv-1* | 1,081 | mScarlet | 0.519 | 0.385 | 0.005 | 0.00E+00 |
| *Y48G10A.6* | 1,316 | mScarlet | 4.48 | 0.987 | 0.01 | 0.00E+00 |
| *F15G9.1* | 5,349 | mScarlet | 2.05 | 0.936 | 0.087 | 6.12E-178 |
| *T25B9.5* | 1,142 | mScarlet | 0.557 | 0.262 | 0 | 0 |
| *K03H9.3* | 4,738 | mScarlet | 0.591 | 0.308 | 0 | 0 |
| PVD | | | | | | |
| *C45B11.6* | 197 | GFP | 0.623 | 0.391 | 0.003 | 1.73E-22 |
| *T05C12.11* | 1,869 | GFP | 0.741 | 0.522 | 0.01 | 1.46E-128 |
| *Y51H7C.3* | 1,010 | GFP | 0.888 | 0.435 | 0.03 | 9.94E-27 |
| *T03F6.4* | 5,788 | GFP | 1.09 | 0.652 | 0.017 | 1.27E-117 |
| *shw-3* | 7,945 | GFP | 0.829 | 0.522 | 0.021 | 1.98E-60 |
| *F21F3.7* | 3,198 | GFP | 1.09 | 0.652 | 0.017 | 8.94E-44 |
| *T14B1.1* | 7,124 | GFP | 0.703 | 0.522 | 0.044 | 1.60E-25 |
| *ver-4* | 3,163 | GFP | 0.447 | 0.261 | 0.001 | 0 |
| *F31A9.2* | 2,242 | GFP | 1.43 | 0.29 | 0 | 0 |
| *Y57G11C.39* | 1,726 | GFP | 0.326 | 0.194 | 0 | 0 |
| *srab-9* | 1,803 | GFP | 0.361 | 0.177 | 0 | 0 |

**Supplementary Table 1. Putative FLP- and PVD-enriched promoters.**
List of putative FLP (top) and PVD (bottom) enriched genes from the CeNGEN database that was publically available in 2020 (Taylor et al., 2021). Genes were chosen based on a high percentage of cells expressing the gene in group 1 ("1%") and a low percentage of cells expressing the gene in group 2 ("2%"). These 25 promoters, defined here as the entire intergenic region between the gene's start codon and the last base pair of the preceding gene's 3' UTR ("Length (bp)"), were used to drive expression of either mScarlet in FLP, or GFPnovo2 in PVD. Priority was given to genes that were shown as enriched in one neuron but not the other, as indicated by the "Enriched Genes by cell type", except in a few cases where the $\log_2$ fold change for a particular gene, such as *Y48G10A.6* and *F15G9.1*, was high.

### Supplementary Table 2: Strains

| Figure | Strain | Genotype | Description |
|---|---|---|---|
| 1a | TV27020 | *wyEx10338* | wyEx10338 = pY48G10A.6::mScarlet + odr-1 GFP |
| 1c, e, f | TV27427 | *wyIs959 IV (“WT”)* | wyIs959 = ser2prom3::ZIF-1 + pY48G10A.6::zf1::mScarlet + ser2prom3::myr-GFP |
| 2a-f<br>4a | TV27427 | *wyIs959 IV (“WT”)* | wyIs959 = ser2prom3::ZIF-1 + pY48G10A.6::zf1::mScarlet + ser2prom3::myr-GFP |
| 2a, b | TV27791 | *mig-14(ga62) II ; wyIs959* | mig-14 (ga62) = loss of function point mutation |
| 2d-f | TV27787 | *cwn-1(ok546) II; wyIs959* | cwn-1 (ok546) = loss of function deletion (787 bp deletion, intron 1 to exon 4) |
| 2d-f<br>3f, g, S2d | TV27788 | *cwn-2(ok895) IV; wyIs957* | cwn-2 (ok895) = loss of function deletion (906 bp deletion, exon 2 to intron 5)<br>wyIs957 = ser2prom3::ZIF-1 + pY48G10A.6::zf1::mScarlet + ser2prom3::myr-GFP |
| 2d | TV27789 | *egl-20(n585) IV; wyIs958* | egl-20 (n585) = loss of function point mutation G295A (C99S) |
| 2d | TV27790 | *lin-44(n1792) I; wyIs959* | lin-44 (n1792) = early stop codon (W100stop) |
| 3a, b, e | TV29053 | *wy1885; glo-1(zu391) X; wyEx10678* | wy1885 = cwn-2::mNeonGreen endogenous C-terminal knockin<br>glo-1 (zu391) X = G to A in 3’ splice site, exon 5 (results in loss of gut granules)<br>wyEx10678 = pY48G10A.6::mScarlet |
| 3b | TV29197 | *glo-1(zu391) X; wyEx10678* | |
| 3c | TV29175 | *wy1885; glo-1(zu391) X; mig-14(ga62) II* | |
| 3d | TV28074 | *wy1717* | wy1717 = CRISPR GFPnovo2 insertion after the C-terminus of cwn-2 |
| 3f | TV28189 | *cwn-2(ok895) IV; wyIs957; wyEx10558* | wyEx10558 = pdpy-7::cwn-2 hypodermis cell rescue + Punc-122::RFP |
| 3f | TV28391 | *cwn-2(ok895) IV; wyIs957; wyEx10558* | wyEx10558 = pdpy-7::cwn-2 hypodermis cell rescue + Punc-122::RFP |
| 3g | TV29199 | *cwn-2(ok895) IV; wyEx10695* | wyEx10695 = plin-44::cwn-2 & unc-122::RFP |
| 3g | TV29198 | *cwn-2(ok895) IV; wyEx10672* | wyEx10672 = plin-44::cwn-2 & unc-122::RFP |
| 3h | TV? | *cwn-2(ok895) IV; glo-1(zu391) X; wyEx?* | wyEx? = pdpy-7::cwn-2::mNG + pY48G10A.6::mScarlet |
| 3h | TV?? | *cwn-2(ok895) IV; glo-1(zu391) X; wyEx??* | wyEx?? = plin-44::cwn-2::mNG + pY48G10A.6::mScarlet |
| 4a-g | TV28078 | *lin-17(n671) I; wyIs959 IV* | lin-17 (n671) = early stop codon (ochre: C1345T) |
| 4a | TV28079 | *mig-1(e1787) I; wyIs959 IV* | mig-1 (e1787) = early stop codon (ochre: Q277*) |
| 4a | TV28080 | *cfz-2(ok1201) V; wyIs959 IV* | cfz-2 (ok1201) = loss of function deletion (1175 bp deletion (1645 to 2819)) |
| 4c, d, g | TV28392 | *lin-17(n671) I; wyIs959 IV; wyEx10583* | wyEx10583 = pY48G10A.6::lin-17 cDNA (PVD + FLP rescue) |
| 4c, d | TV28400 | *lin-17(n671) I; wyIs959 IV; wyEx10583* | wyEx10583 = pY48G10A.6::lin-17 cDNA (PVD + FLP rescue) |
| 4c, e, g | TV28426 | *lin-17(n671) I; wyIs959 IV; wyEx10593* | wyEx10593 = pY48G10A.6::zf-lin-17 cDNA (FLP rescue) + Punc-122::RFP |
| 4c, e | TV28530 | *lin-17(n671) I; wyIs959 IV; wyEx10593* | wyEx10593 = pY48G10A.6::zf-lin-17 cDNA (FLP rescue) + Punc-122::RFP |

**Supplementary Table 2: Strains** (*Continued*)

| | | | |
|---|---|---|---|
| 4c, f, g | TV28425 | *lin-17(n671) I; wyIs959 IV; wyEx10592* | wyEx10592 = ser2prom3::lin-17 cDNA (PVD +OLL) rescue + Punc-122::RFP |
| 4c, f | TV28434 | *lin-17(n671) I; wyIs959 IV; wyEx10592* | wyEx10592 = ser2prom3::lin-17 cDNA (PVD +OLL) rescue + Punc-122::RFP |
| 5a-g | TV29053 | *wy1885; glo-1(zu391) X; wyEx10678* | wy1885 = cwn-2::mNeonGreen endogenous C-terminal knockin<br>glo-1 (zu391) X = G to A in 3' splice site, exon 5 (results in loss of gut granules)<br>wyEx10678 = pY48G10A.6::mScarlet |
| S1 | TV27020 | *wyEx10338* | wyEx10338 = pY48G10A.6::mScarlet + odr-1 GFP |
| S2a | TV27130 | *dma-1(tm5159) I; wyEx10338* | dma-1 (tm5159) = loss of function deletion (672 bp) |
| S2b | TV27131 | *sax-7(nj48) IV; wyEx10338* | sax-7 (nj48) = loss of function deletion (568 bp) |
| S2c | TV27541 | *Y48G10A.6(wy1643) I; wyIs959* | Y48G10A.6 (wy1643) = loss of function deletion of entire coding region |
| S3 | TV29053 | *wy1885; glo-1(zu391) X; wyEx10678* | |
| S4a | TV28433 | *wy1778* | wy1778 = CRISPR GFPnovo2 insertion after the C-terminus of lin-17 |
| S4b, c | TV27427 | *wyIs959 IV ("WT")* | wyIs959 = ser2prom3::ZIF-1 + pY48G10A.6::zf1::mScarlet + ser2prom3::myr-GFP |
| S4b, c | TV28078 | *lin-17(n671) I; wyIs959 IV* | lin-17 (n671) = early stop codon (ochre: C1345T) |
| S4b, c | TV28392 | *lin-17(n671) I; wyIs959 IV; wyEx10583* | wyEx10583 = pY48G10A.6::lin-17 cDNA (PVD + FLP rescue) |
